# Local climate variability, phenology and morphological traits of the long distance migrants Savi’s warbler *Locustella luscinioides* and Sedge warbler *Acrocephalus schoenobaenus* in reedbeds of a man-made wetland of SE Iberia

**DOI:** 10.1101/2024.05.24.595692

**Authors:** Ignacio García Peiró

## Abstract

As a consequence of climatic variability in the northern hemisphere, the Mediterranean region is experiencing the most pronounced effects of rising temperatures and decreasing precipitation. This has a multitude of effects on bird migration, with particular relevance to migratory passerines associated with wetlands, whose area has been in decline in this region since the last century. In order to relate phenology to morphology and climate in two species of trans-Saharan migrants closely associated with reedbeds, this study analysed the relationships between Savi’s warblers *Locustella luscinioides* and Sedge warblers *Acrocephalus schoenobaenus* abundances, local climatology and morphological traits in an inland coastal artificial wetland in southeastern Iberia over a 12-year period. The climatic variability observed in this wetland was reflected in a negative trend between local temperatures and the year, and a positive trend with annual precipitation. This was confirmed in subsequent series. The abundance of Savi’s warbler increased adaptively with the year, while that of the Sedge warbler decreased non-adaptively, although neither change was statistically significant. A two-year delay was observed in the interannual phenology peak of the Savi’s warbler relative to the Sedge warbler. The monthly abundances of Savi’s warblers exhibited a significant positive correlation with intra-annual temperatures, explaining approximately half of the intra-annual phenology. No morphological trait could be identified as an explanatory factor for these trends, as no significant correlation with year was detected. Consequently, a coincidence with the morphological traits of both species associated with global climate change could not be established, which supports the hypothesis of migratory bird mismatch in the southeastern Iberia. In a future scenario in the eastern fringe of the Iberian Peninsula, an increase in the abundance of some trans-Saharan migrants, such as the Savi’s warbler, is to be expected as a consequence of climatic improvement, in particular rising temperatures. Further studies are required to ascertain whether this phenomenon occurs in other trans-Saharan migrants in other localities in the west.

## Introduction

As a result of climate change and/or weather variability in the northern hemisphere, the Mediterranean region is being the most affected by increasing temperatures and decreasing precipitation (Sanz 2002, Xoplaki 2002, Lionello *et al*. 2006). Migratory birds respond to these effects in a variety of ways (see reviews in Lack 1960, Crick 2004). The relationship of phenology and morphological traits in relation to climate change has been subject of multiple studies for recent decades, and the patterns of response remain unclear because of its complexity which affects the phenological processes. This is due to the underlying problems inherent to sexes, ages, and departure times and distance (MacMynowski and Root 2007, Verhoeven *et al*. 2022) so may compromise the conclusions due to the relationships may be non-linear (Saleski *et al*. 2013, Pearce-Higgins *et al*. 2015) despite morphological traits (e.g. body size) that reflect relationships in response to climate change can be non-linear (Gardner *et al*. 2014). In addition they have a summed effect of constraining the migration life-history stage of natural populations (Zimin *et al*. 2023) and various studies reflect that the phenology may match climate (Renfrew *et al*. 2013), mismatch (Saino *et al*. 2011) or both (Stenseth and Mysterud 2002), may be insufficient (Radchuk *et al*. 2009) and inconclusive (Neto *et al*. 2009) and then may compromise the phenotypic plasticity (Charmantier *et al*. 2008).

Population sizes that are associated with climate change are decreasing (Møller & Merila 2004, Miller-Rushing *et al*. 2008, Lindén *et al*. 2017) and these may be due to potential problems inherent to the interpretation of long-term data and in particular those that offer an explanation as responses to climate change, already claimed by Møller and Merilä (2004). For example, Knudsen *et al* (2007) indicate the existence of inherent problems about how to construct robust frameworks of the complete phenology, with proper measures of centrality and dispersal, without analyzing dates on arrivals and departures because they may be erroneous (Sparks *et al*. 2007) or inconsistent (Wilson 2009). Møller and Merilä (2004) suggest that use of medians, medians and modes should be a good parametric predictor in analysis contrasting with climate change but these data are still few studied.

Few studies have been made trying to seek what variables match better the migration periods and annual phenology of long-distance migrants. The absence weather study variables (e.g. relative humidity, cloud cover, sun radiation, wind speed and direction) was a key in the past and yet results a handicap for the treatment of the responses of bird’s phenology to climate change despite of some studies make use of these datasets and put hands on (Arizaga *et al*. 2011, Haest *et al*. 2019, 2020, Bozó *et al*. 2021) but others consider some variables traditionally used for analyzing the phenological responses to climate change are already undervalued or give more importance to other environmental variables when confronted with phenology (Gordo 2007) so the variable choice is crucial because can affect the species vulnerability in relation to climate change (Oswald and Conradie 2023) although the inclusion of multiple climatic variables is already a fact (Remisiewicz and Underhill 2020).

Confronting the phenology and morphological traits with climatic variables is important due to the impact of climate change on the population dynamics and its body components (Cotton, 2003, Charmantier *et al*. 2008, Kovács *et al*. 2012). In this sense, is observed a hugh inter-(Rubbolini *et al*. 2007) and intra-specific variability (MacMynowski and Root 2007) on which phenotypic plasticity plays an important role (Crick 2004) in an adaptive (Charmantier *et al*. 2008), maladaptive (Remacha 2020) or both forms (Radchuk *et al*. 2019).

The autumn migration of the Sedge warbler *Acrocephalus schoenobaenus* (hereafter ASCH) is well studied in the past (see Bibby *et al*. 1976, Bibby & Green 1981, Bibby and Green 1983, Ormerod 1990). It is showed to be strongly age and sex-dependent (Péron *et al*. 2007, Miholcsa *et al*. 2009, Jakubas and Jakubas 2010, Kovács *et al*. 2012) and highly dependent on the winter survival in sub-saharan Africa (Peach *et al*. 1981; Baillie and Peach 1992) that in turn is dependent on weather conditions there (Haest *et al*. 2018, 2020). From the viewpoint of the optimality of migration, ASCH behaves as time-selective migrant because of the fuel deposition rates are not directly related with with departure rates, at least partially (Bayly 2007) and the stay sites are highly dependent of the unpredictable food availability based in a thermo-dependent prey as the plum-reed aphids (Bibby and Green 1976, 1981, Peach *et al*. 1981). The availability of this prey is highly variable both spatially and temporally (Nartshuk 1996) and clearly susceptible to weather conditions (Chertnesov 1998, Chernetsov and Manukyan 2000), which, contrarily to other *Acrocephalidae* (Arizaga *et al*. 2011, 2020) seem not to affect the departure decisions at least partially (Bayly 2007) although lately it has been showed that inbreeding depression may seen affected by environmental conditions in the Aquatic warbler *Acrocephalus paludicola* (Kubacka *et al*. 2004). The spring migration of ASCH appears to be strongly dependent on weather conditions in African Sahel (Foppen *et al*. 1999, Haest *et al*. 2020 and references therein) and on climatic conditions at European stop-over sites (Hakka *et al*. 2011).

The autumn migration of the Savi’s warbler *Locustella luscinoides* (hereafter LL) seems to be linked to moult processes in breeding and wintering areas, which impose a substantial constraint in space and time in the annual cycle (Neto *et al*. 2006, Neto and Gosler 2006). LL undergo a complex post-juvenile and adult post-breeding moult termed “*filling moult*” (Neto and Gosler 2006, Kelemen *et al*. 2000), and its extension is fledging date (Svensson 1992) and body condition dependent (Neto an Gosler 2010, Kulaszewicz and Jakubas 2015). The complete moult occurs at sub-Saharan winter quarters (Neto *et al*. 2006). The autumn migration of Savi’s warbler is clearly age-dependent where adults are time-selected migrants and juveniles time-minimizing migrants (Neto *et al*. 2008). Adults, in contraposition to juveniles, migrate later and materialize short-time stopovers where they gain greater fuel deposition to reach longer distances (Neto *et al*. 2008). From the feeding behavioral viewpoint LL is a “*foraging-time limited migrant*” limited by food availability (Lindström 1991, Neto *et al*. 2008).

Although many studies have been carried out in recent decades investigating the phenology of Sylviidae trans-Saharan migrants specialized in reed marshes and swamps in relation to climate change and morphological traits, these studies remain already scarce (e.g. Kaňuščák *et al*. 2004; Péron *et al*. 2007, Halupka *et al*. 2008). For this reason, I try to infer some aspects of the phenology of two long-distance strict migrants that stopover in the southeastern corner of Iberia and its relationships under climate conditions, setting future prospects in the study of Mediterranean wetlands which have undergone a huge decline in surface from the past century (Perennu *et al*. 2012) and they are of high priority for some Eurasian threatened long-distance reedbed warblers.

Based on the earlier assumptions and framework, I analyzed a 12-year climatic dataset (1992-2003) from an inland coastal wetland in the southwestern Mediterranean (SE Iberia) and compared it with a database of the ringing captures of a tiny population of the two species. On doing so I use some components of phenology and confront them with climate and morphology, setting and trying to solve the following hypothesis: (1) Do the trends in the climate variables studied and local scenarios not differ from the prospects observed at Mediterranean scale or alternatively exist differences?, (2) are there no differences in measures of centrality and dispersal that defines the intra- and inter-annual phenology, as well as in the fluctuations of the body size index and body condition index of the migrants studied or by the contrary are found different patterns and also its relationships ?, (3) can this possible absence of variability mismatch the climate inter- and intra-annually and then their responses to be maladaptive or by the contrary matching being adaptive ?, (4) what explanations can be offered to confirm or reject the earlier hypothesis? and finally, (5) what conservation purposes can be extracted from the results for reedbeds and hence for the high priority specialists’ warblers associated with wetlands which have undergone a huge decline in surface area from the past century ?.

## Material and Methods

### The study area

The study area belongs to the Thermo-Mediterranean-Iberian bioclimatic sector according to Rivas-Martinez (1983). In this context, the south-eastern Iberian plateau conforms a transitional boundary with the arid and semi-arid regions of the provinces of Alicante (38°20’N 0°28’W) and Murcia (37°59’N 1°7’W), covering an area of approximately 108,000 Ha. A lowland plateau in the middle south of the province of Alicante, formed mainly by the catchment areas of the River Segura, has overgrown a vast marshland of approximately 20,000-40,000 ha. At the beginning of the XX century, an artificial inland reservoir was created (*Navarro, 1988*). This swamp was declared Ramsar site number 14 in 1971 under the name of "El Hondo Swamp" (Bernués 1989, SEOBirdlife 2012) and in 1988 as "El Hondo Natural Park": 38°10’N 00°44’W (GVA 1989) due to its international importance for shorebirds and waterbirds. The community of passerines is rich in *Sylviidae* Afrotropical migrants with the Eurasian Reed warbler *Acrocephalus scirpaceus* and the Great Reed warbler *Acrocephalus arundinaceus* conforming the bulk of the breeding community (Peiró 2006). Currently, it is immersed in a fragmented agriculture landscape as an output-input of a reservoir and freshwater is supplied by channels of the Tajo-Segura Iberia river connections. The Park is 15 km from the Mediterranean sea and 360 km north from the north coast of Argelia (Figure 1).

**Figure 1.**
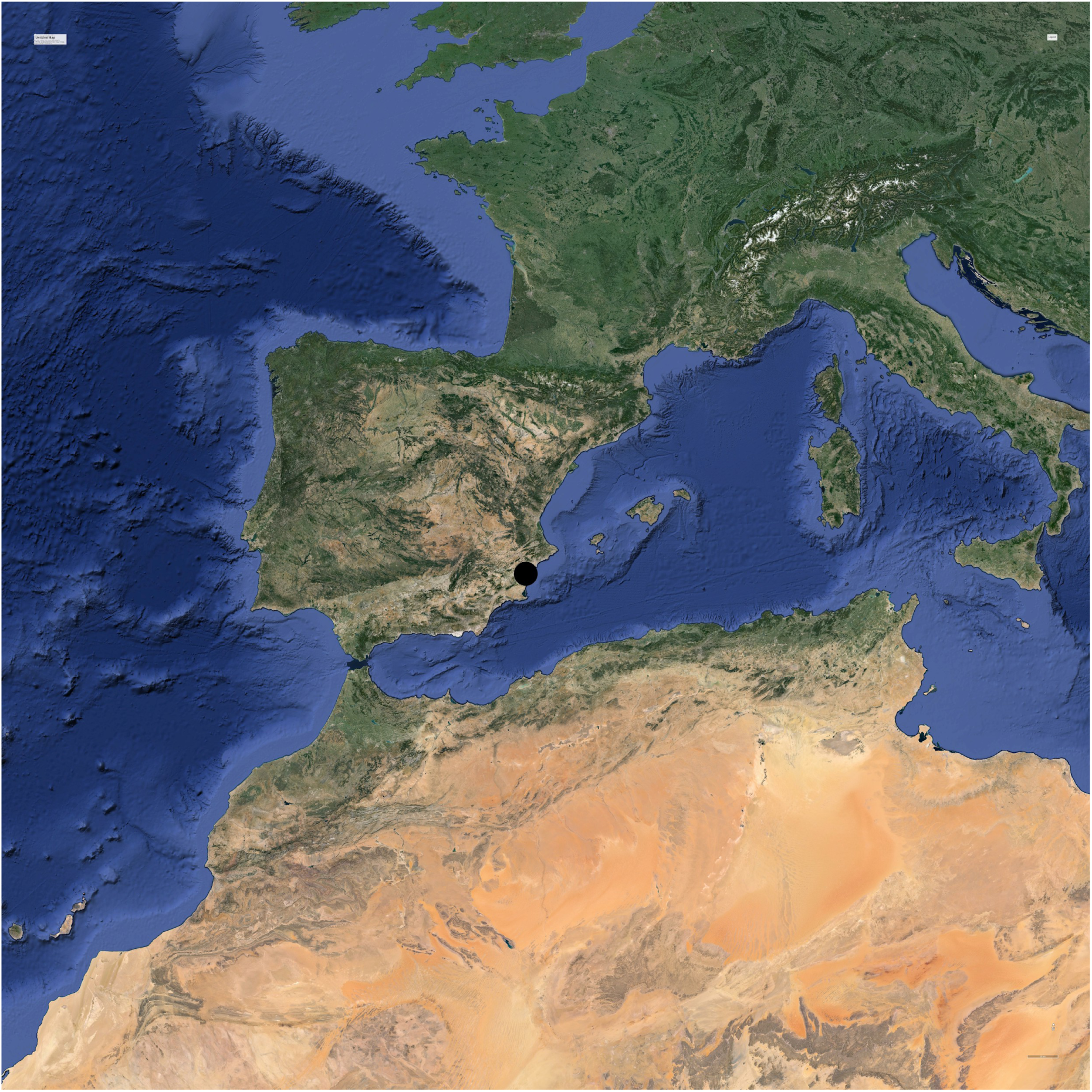
Ubication of the site “El Hondo Natural Park” in southeastern corner of Iberia.

### Datasets Treatment

#### Climatic data

In order to give an overall climatic sketch of the site I have collated a dataset spanning 12-years (1992-2003) in a meteorological station inside “*Estación Experimental Agraria of Elche-Generalitat Valenciana*” (EEA-IVIA) 11 km NE from the Park at 38°14’N 1°25’W and 62 meters over sea level. Original climatic dataset was, (1) daily maximum and minimum temperatures (°C) and (2) monthly total rainfall (mm). These variables were transformed in the following climatic proxies: 1) Mean monthly temperature (°C) from 1992 to 2003, 2) annual summed total precipitation (mm) of spring (February-May) and autumn (July-October) respectively from 1992 to 2003, 3) mean annual temperature in 1992-2003 and 1991-2020 4) overall annual precipitation in 1992-2003 and 1991-2020. These data were chosen as a trade-off with local arrival dates of both warblers and local overall scenario of the climate change. Original datasets were available online in http://riegos.ivia.es/datos-meteorologicos.

#### Population data

In support of climatic data, some population parameters such as phenology and body components of ringed birds were used. Most ringing visits were carried out in the morning, when the bird’s activity is higher (Brunil *et al*. 2014, Farina *et al*. 2015) but in some circumstances were conducted in the afternoon, so this might have influenced the probability of taking a more or less species in some months and years of research. However, dawn and evening sampling is found suitable for estimating the majority of reed passerines species (Trnka *et al*. 2006). For trapping birds, I standardized the sampling effort per season and year. I only used the trapping effort of those months where the birds were trapped, considering the spring season (from February to May) and autumn season (from July to October). I used a variable number of ringing sessions employing a mean session of 73.9 m net·hour + 72.5 in spring and 74.9 m net·hour + 73.6 in autumn (t = −0.071, P = 0.948). The annual effort was variable and was considered summing spring and autumn periods, with a mean of 105.25 m net·hour + 99.67 per year. I calculated the accuracy of the samples by means of the variation coefficient. Other authors studying effectivity of mist netting of marsh passerines (Poulin *et al*. 2000) indicate 250 m net·hour each session is sufficient to detect the population of reed passerines. Considering this, the mean annual effort per session was 42 % of the optimal per session. The number of birds ringed was 18 for LL and 17 for ASCH. Most LL ringed were juveniles (89%) while the proportions of juveniles and adults in the sample of ASCH was similar (38% and 42%, respectively). One juvenile bird ringed on 24th July 1993 and recovered 28th August 1993 showed that probably a tiny part of the sample were of local birds despite that in the Spanish Breeding census (1998-2002) it wasn’t recorded as breeder (López *et al*. 2003) but was detected posteriorly (2008-2014: Vera 2022).

Due to the earlier, I estimated the minimum new sample size a priori for LL and ASCH using the G-Power sample size program (Faul *et al*. 2007) and using the original monthly captures. This was done by calculating the differences between two independent arithmetic means and standard deviations (LL = 4.5000 ± 3.3166; N = 18; ASCH = 2.4286 ± 2.2951; N = 17), with an effect size of 0.7532, allocation ratio of 0.9444, overall standard deviation of 2.7502, a power of 0.95 and a significance level of 0.05. This test was repeated three times. The new number of individuals estimated under these conditions was LL = 40 and ASCH = 38. Due to the low estimated sample size, I increased the power to 0.99 with a significance level of 0.01. The new number of estimated samples was LL = 80 and ASCH = 76. The overall increase was significant (Chi-squared test: X_1_^2^= 26.00; p < 0.0001).

To define phenology, I firstly calculated, as centrality measures, the arithmetic mean, modes and median of the annual and monthly frequency distribution of birds ringed in 1992-2003 and published in Peiró *et al* (2005) but subsequently reckoned using my own ringing database. The modes were used to detect skew in the distributions and were tested by means of the Excess-Mass test (Müller & Sawitzki 1991, Ameijeiras-Alonso *et al*. 2019). The medians were used to detect shifts in the distributions and were tested by Willcoxon Signed-rank tests (Sokal & Rohlf 1981) The variation coefficient was used to measure dispersal of the distributions. The original ringing captures of both warblers were confronted to the climatic dataset and the mean morphological indexes (Figure 2).

#### Morphological traits

I metal-ringed each bird (Ministery of environment-MADRID-SPAIN and identified by plumage features (Peterson et al. 1977) and aged as juveniles or adults according to Svensson (1992). Afterwards it was recorded the body mass with a ©Pesola spring balance (to the nearest 0.1 g), the maximum right wing length (to the nearest 0.5 mm), the bill-length to the skull (to the nearest 0.1mm), the right tarsus-length (to the nearest 0.1 mm) to the last scales with a © Camblab caliper, according to Svensson (1992). Fat score was recorded as a visual index (scale 0-5, Helms and Drury, 1960). It was not possible to record the repeatability of measures, so it was not possible to discern among measurement error and ontogenetic variation (Jones and German 2005). Instead was calculated the variation coefficient (Table 1). The combination of the earlier set of measures launched two body traits: (1) Body structural size Index (BSI). Despite the underlying problems of interpretation of size of birds on PCA (Rising and Somers 1989, Freeman and Jackson 1990, Piersma and Davidson 1991, Schamber *et al*. 2009) BSI was considered as residuals of the linear regression of tarsus length on wIng-l length because this kept a good determination coefficient on the correlation with bill-length (LL: Wing-length = 1.468·Tarsus-length + 38.670; r^2^ = 0.107, P = 0.2545.; ASCH: Wing-length = 1.723·Tarsus-length + 30.890; r^2^ = 0.199, P = 0.1088), (2) Body Condition Index (BCI) was considered as the residuals of a linear regression of fat on wing-length (LL: Body-weight = 0.859·Fat + 12.322, r^2^ = P < 0.00; ASCH: Body-weight = 0.307·Fat + 10.789, r^2^ =, P = 0.057) being a good estimator of body condition (see Schamber *et al*. 2009; Labocha and Hayes 2012). Body weight is a measure of body mass that is plastic trait highly variable intra- and inter-annually and highly dependent on fat reserves on migration (Robson & Barriocanal 2008, Neto *et al*. 2008) and as a proxy of physiological processes through immunocompetence signal (Møller *et al*. 1998, Møller and Petrie 2002), stress during breeding (Neto and Gosler 2010) and as fitness signal (Fenoglio *et al*. 2002).

**Table 1.**
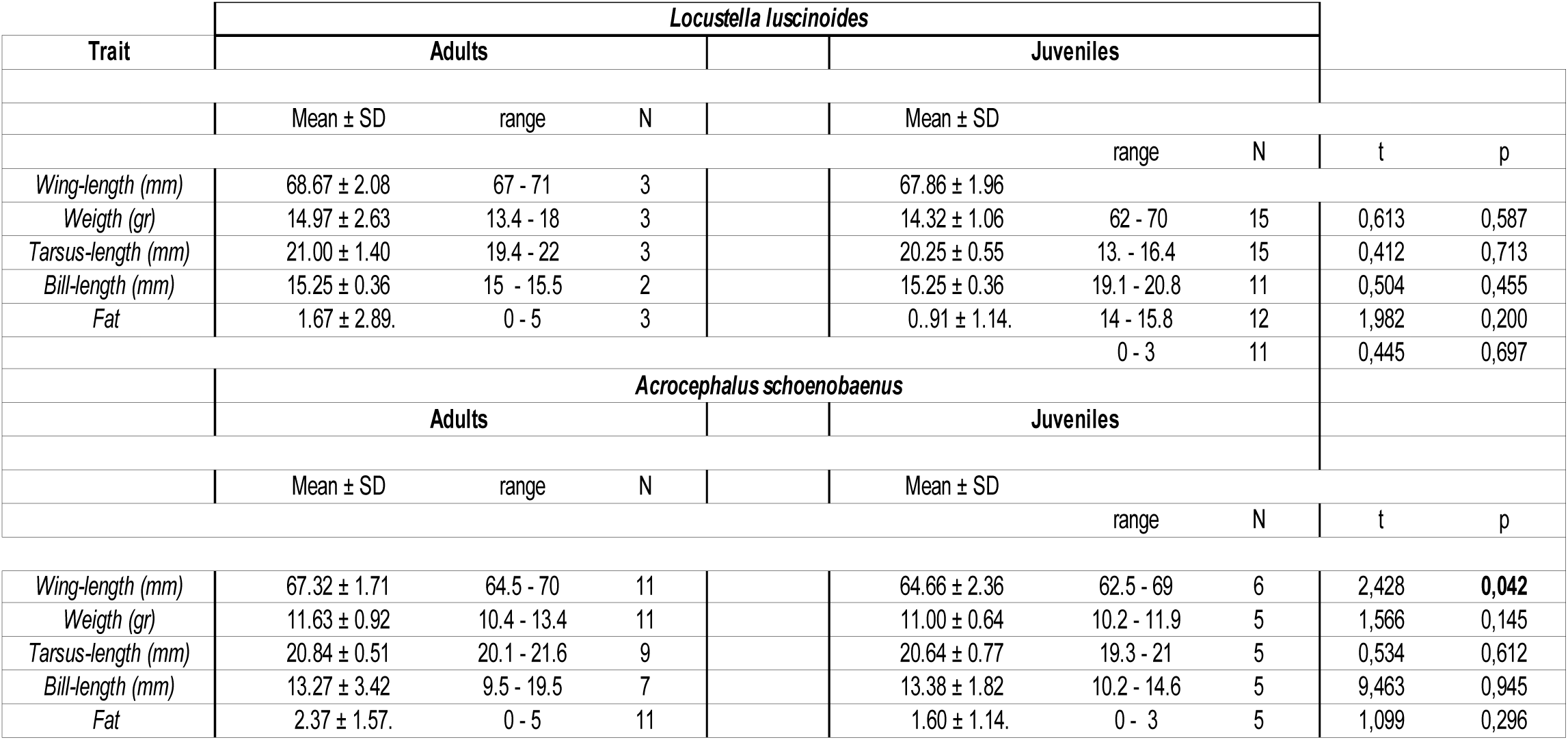
Biometrics of LL and AS in “El Hondo Natural Park”.

**Table 2.** *General Linearized Models* (GLM) for the intra-annual phenology and inter-annual phenology of LL and ASC. Best AIC and BIC models are selected. BSI was obtained as the mean annual residuals of the linear regression of tarsus-length on bill-length. BCI was considered as the mean monthly residuals of the linear regression of fat on wing-length. Monthy mean temperature (°C) and total monthly rainfall (mm) are depicted also as independent-predictive variables in the linear regressions. Significant p values (p < 0.05) are in bold.

| *Acrocephalus schoenobaenus* |  |  |  |  |  |  |  |  |  |  |
| --- | --- | --- | --- | --- | --- | --- | --- | --- | --- | --- |
| Intrannual |  |  |  |  |  |  |  |  |  |  |
| VARIABLE | b | SE | t | p | AIC | BIC | F | df | P | r2 adj |
| INTERCEPT | 1,473 | 0,640 | 2,301 | **0,044** | 56,614 | 58,069 | 0,256 | 1,10 | 0,624 | -0,073 |
| MONTHLY BCI | -0,809 | 1,559 | -0,506 | 0,624 |  |  |  |  |  |  |
| Interannual |  |  |  |  |  |  |  |  |  |  |
| VARIABLE | b | SE | t | p | AIC | BIC | F | df | P | r2 adj |
| INTERCEPT | 2,574 | 1,492 | 1,726 | 0,115 | 49,830 | 51,285 | 0,670 | 1,10 | 0,432 | -0,031 |
| ANNUAL RAINFALL | -0,008 | 0,010 | -0,819 | 0,432 |  |  |  |  |  |  |
| *Locustella luscinioides* |  |  |  |  |  |  |  |  |  |  |
| Intrannual |  |  |  |  |  |  |  |  |  |  |
| VARIABLE | b | SE | t | p | AIC | BIC | F | df | P | r2 adj |
| INTERCEPT | -5,810 | 2,328 | -2,496 | **0,032** | 55,185 | 56,640 | 10,550 | 1,10 | **0,009** | 0,465 |
| MONTHLY TEMPERATURE | 0,379 | 0,117 | 9,248 | **0,009** |  |  |  |  |  |  |
| Interannual |  |  |  |  |  |  |  |  |  |  |
| VARIABLE | b | SE | t | p | AIC | BIC | F | df | P | r2 adj |
| INTERCEPT | 1,624 | 0,736 | 2,208 | 0,052 | 59,951 | 61,405 | 0,955 | 1,100 | 0,352 | -0,004 |
| ANNUAL BSI | 0,306 | 0,313 | 0,352 | 0,952 |  |  |  |  |  |  |

#### Statistical treatment

##### Climatic data

Climate data did meet the normality (Monthly mean temperature: Shapiro-Wilk, W = 0.971, P = 0.1985; Annual mean temperature: W = 0.638, P = 0.0546; Monthly precipitation: W = 0.940, P = 0.501; Annual precipitation: W = 0.969, P = 0.907), so these variables were considered as parametric. Simple linear regressions between years and climate variables were used to detect possible significant trends. Pearson’s correlations with t-test were used to test these trends.

##### Morphological traits

Some biometrical measures didn’t meet normality (LL: wing length: W= 0.867, P = 0.013, N = 19; body mass, W = 0.848, P = 0.009 N = 18; tarsus-length: W = 0.952, P = 0.586, N = 14) but met for ASCH and were considered parametric. Pearson’s correlations were used to settle possible relationships with body components. Means were expressed as standard deviations (SD). Student-t tests were used to compare mean (Table 1).

##### Relationships between inter, intra-annual phenology, climate and morphological traits

I used *Generalized Linear Models* (GLM: McCullagh and Nelder 1983) between the original abundance of each bird (response variable) and annual mean temperatures, monthly mean temperatures, annual overall precipitation, monthly overall precipitacion and Body Size Index and Body Condition Index as independent-predictive variables using the software package “glm” (Marschner 2011) in the R-Ptoject v 4.0.4 statistical program (R Core Team 2021). Four models were constructed step by step using only basic interactions (n = 4) to ensure more statistical power and the best model was chosen as one with lowest *Akaike’s Information Criteria* (AIC) value. In case of similar AICs I used the lowest value of *Bayesian Information Criteria* (BIC) to elucidate the best model.. Statistical significance of models in Anova’s F-tests were considered at P < 0.05.

## Results

### Trends of climatic variables

From 1992 to 2003, El Hondo Natural Park had an average annual temperature of 20.1°C ± 3.0 and an average annual precipitation of 213.0mm ± 67.9. The study period had significantly 34% lower spring temperatures (16.1°C ± 1.1) than autumn (24.5°C ± 1.4) (Wilcoxon-test = 0, P = 0.000) and rained a non-significant 16% more (spring: 76.3 mm ± 36.7; autumn: 63.8 mm ± 36.8; Wilcoxon-test = 84, P = 0.507). There was a non-significant trend towards colder and wetter years during this period (1992-2003: Temperature:-0.061·year + 140.136, F_1,10_ = 0.428, P = 0.5277; Precipitation: 1.428·year −2638.690, F_1,10_ = 0.058, P = 0.815). These trends were similar over series of 30-years (1991-2020: Temperature: −0.009·year + 36.244, F_1,28_ = 0.1685, P = 0.649: Precipitation: 2.816·year - 5399.480, F_1,28_ = 2.438, P = 0.129). Cold spells increased but not significantly along the study period (r = 0,307, t = 1.019, P = 0.3324). This climatic cycle was governed by autumn temperatures and winter rainfall since its adjusted determination coefficients were the highest on the regression with annual values of temperatures (r^2^ = 0.747) and rainfall (r^2^ = 0.282). This scenario represents a colder and wetter spot in the study area. This is conclusive of a great variability of Mediterranean climatic scenario at local scale as depicted by Xoplaki (2002) in Greece at localities of the Iberian levant Mediterranean (Pereira *et al*. 2021).

### Morphological traits

Adults of both warblers got longer measures than juveniles but only reached significance in wing-lengths of ASCH. Bill-length and fat that were contrary a bit greater in juveniles than adults in ASCH, although without reaching significance (Table 1).

### Overall scenario of the phenology of both transaharian migrants in the study area and Iberia

The phenological pattern of ASH in the EH is mainly prenuptial, with a greater number of birds ringed in spring than in autumn (Figure 2a). The estimated monthly pattern is in agreement with the phenological pattern based on the number of recoveries in Spain (r = 0.299, t = 0.994, P = 0.344, N = 94), which hypothesizes that transaharian spring warblers migrate significantly towards eastern Spain, while in autumn they show an opposite trend (Bernis 1963, Cantos 1991), which is specifically tested based in 48 recoveries of ASCH in Spain (data from Cantos 1988) where the number of recoveries in the autumn passage of ASCH is spreaded towards western (22 vs 6; X_1_ ^2^ = 8.33; P = 0.004) while in spring is similar (5 vs 5) due to the lower sample size. Regarding the phenology of LL in EH (Figure 2b) it is mainly autumnal, which is consistent with its pattern in Iberia (Neto *et al*. 2008).

**Figure 2.**
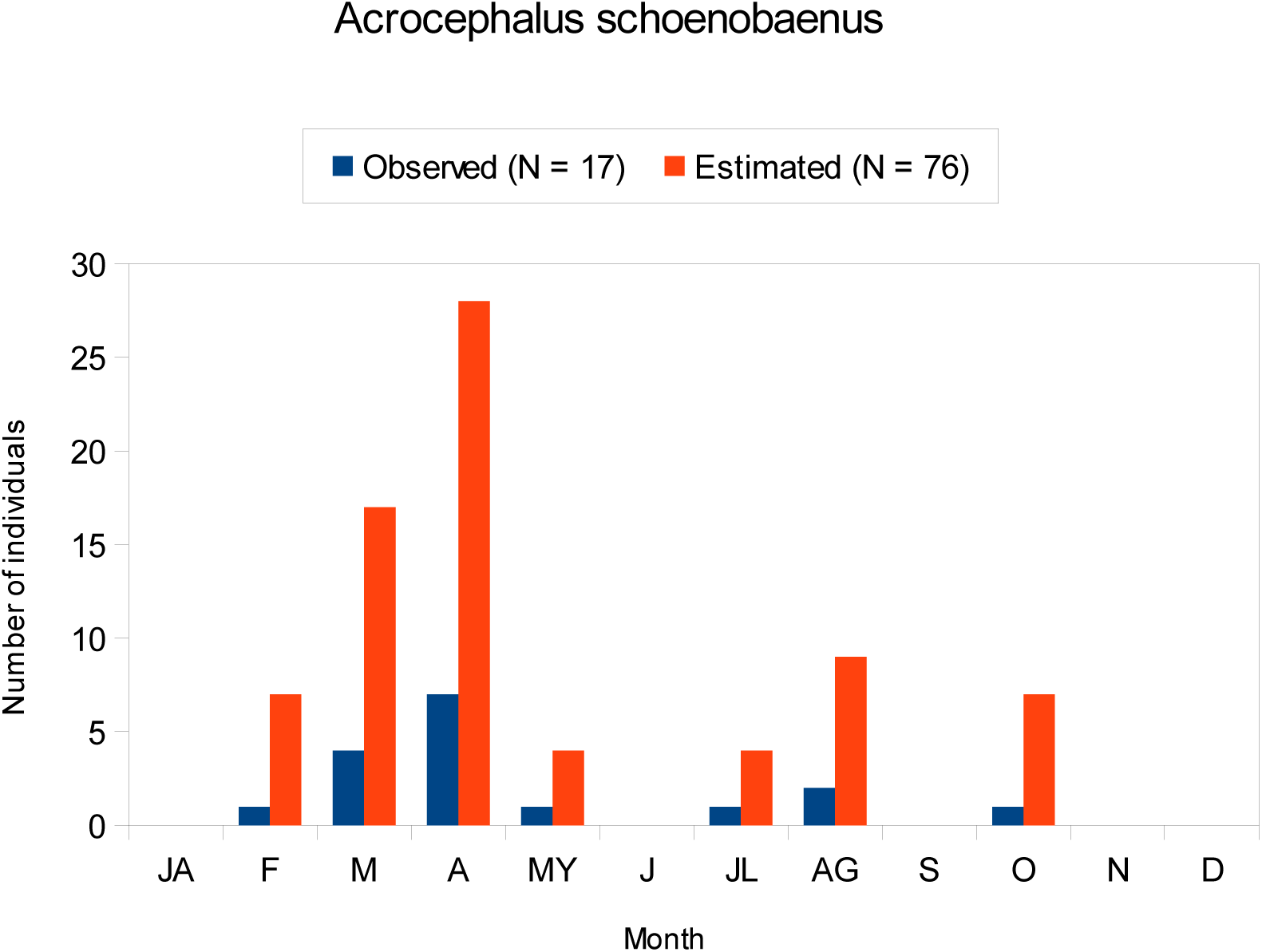
a) ntra-annual phenology of ASCH in “El Hondo Natural Park”. It shows data from the original sample (blue: N =17) and the new estimated sample (red: N = 76). Both samples matched monthly (r = 0.992, t = 24.580, P < 0.0001).

**Figure 2.**
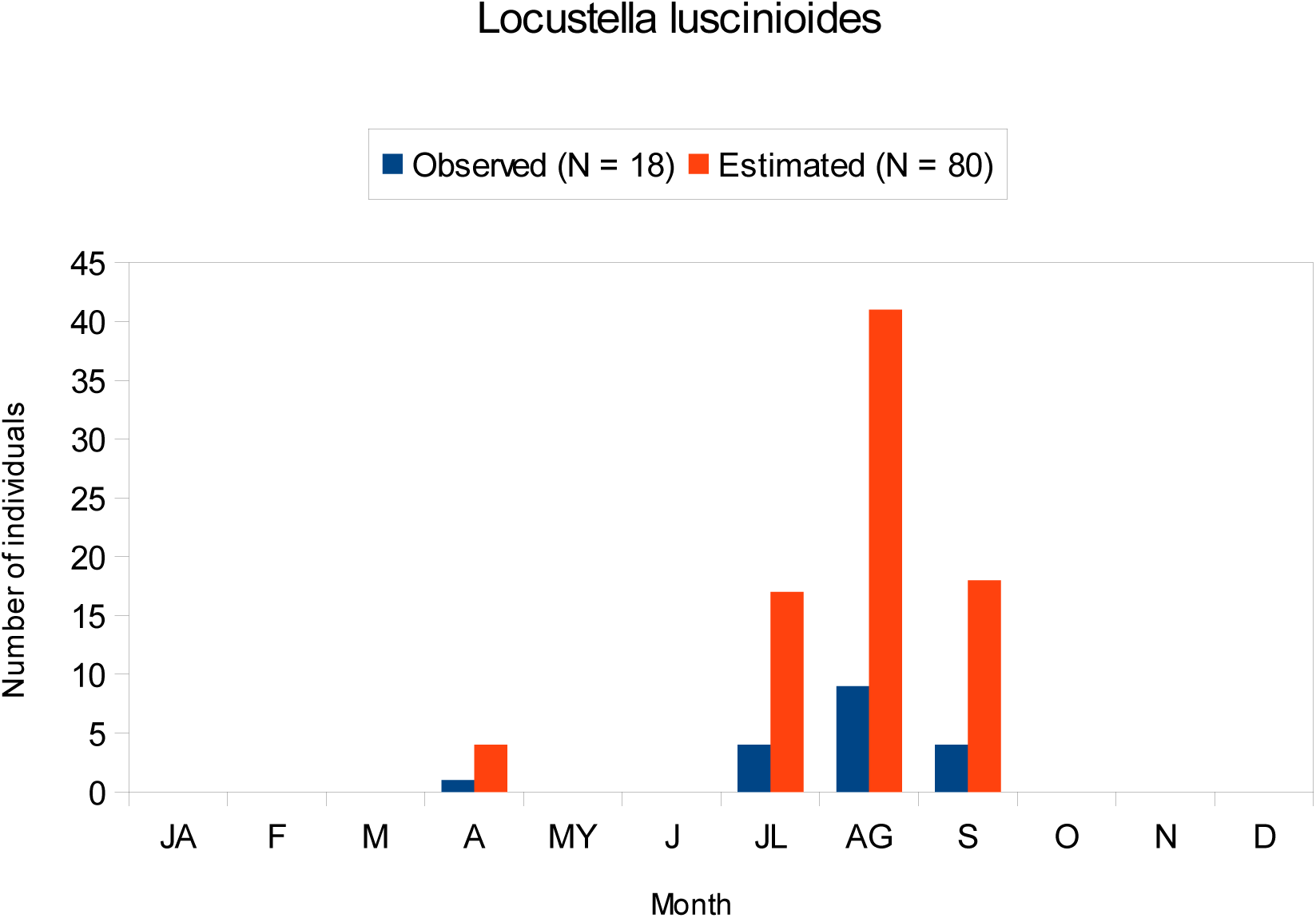
b) Intra-annual phenology of LL in “El Hondo Natural Park”. It shows data from the original sample and the new estimated sample. Both samples matched monthly (r = 0.999, t = 158.63. P = < 0.0001).

### Intra-annual phenology of the transaharian migrants in SE Iberia

The observed sample didn’t meet normality (Shapiro-wilk, LL = 0.622, P = 0.0007; ASCH = 0.708, P = 0.005). The number of individuals per month did not differ among both warblers (t = 0.0573, P = 0.955) but the dispersal differed (LL: CV = 1.89; ASCH: CV = 1.49; Chi-squared: 9.307, P = 0.0023) hence the distribution pattern differed (Figure 2c, d). The phenology of ASH was unimodal (Mode ASCH = 3.598, Excess mass test = 0.079, P = 0.116) and prenuptial, with a greater number of birds ringed in spring than in autumn (Figure 2a) which shows that this warbler migrates significantly in spring in eastern Spain (Cantos 1988), while in autumn they show an opposite trend in similar to the reported by Bernis (1963) for afro-tropical migrants in Iberia. The phenology of LL was unimodal (Mode LL = 8.015, Excess-mass test = 0.067, P = 0.304) and autumnal (Figure 2b), which is consistent with the pattern found by Neto et al (2008) in Portugal. The median of the monthly frequencies distribution for LL was delayed 4 months with respect to ASCH (Median LL = 8; Median ASH = 4; Wilcoxon-test = 5024; P < 0.0001).

**Figure 2.**
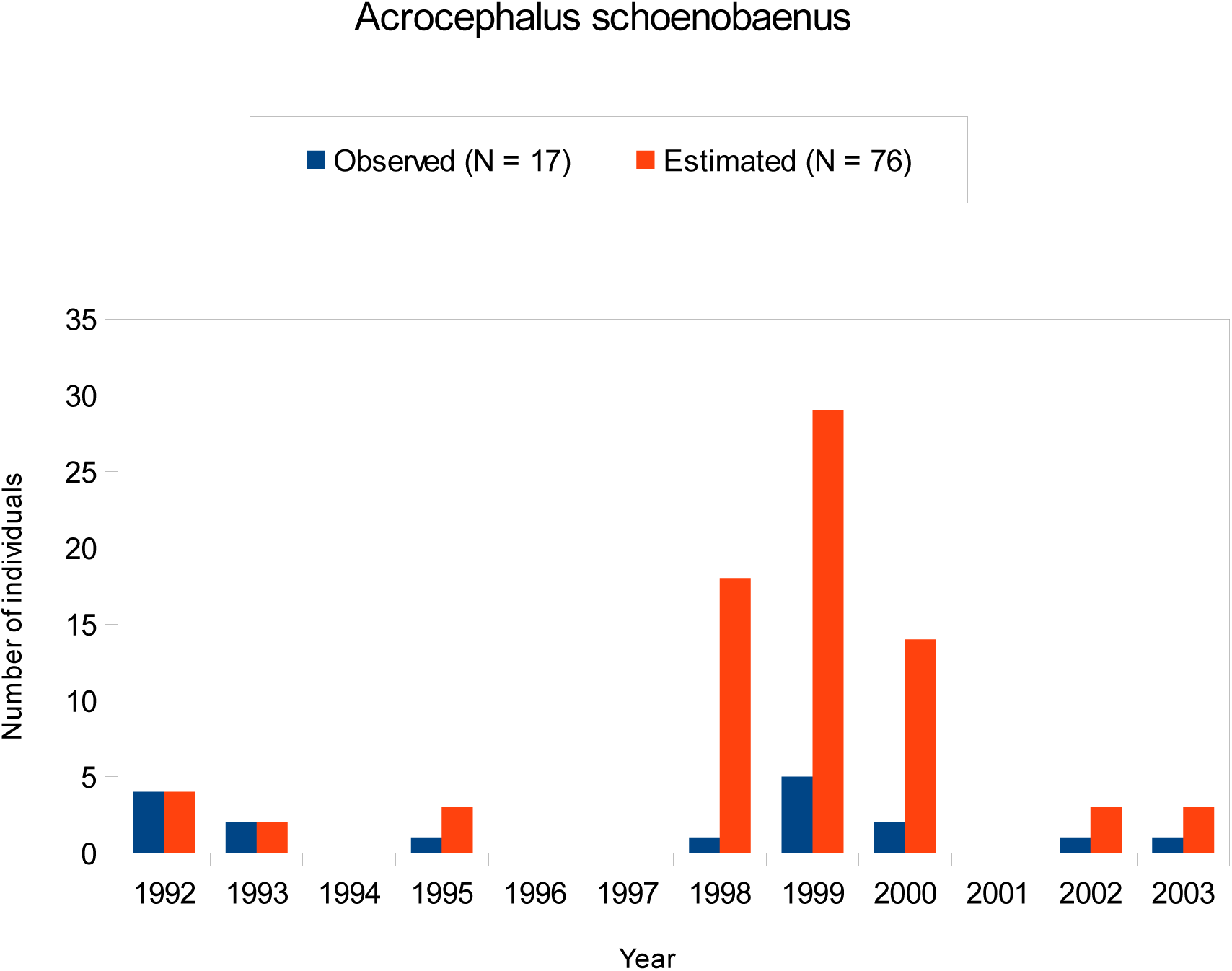
c) Inter-annual phenology of ASCH in “El Hondo Natural Park”. It shows data from the original sample (N = 17) and the new estimated sample (n =76). Original sample (N = 17) and estimated sample (N = 76) matched annually (r = 0.688, t = 2.939 P = 0.0148).

**Figure 2.**
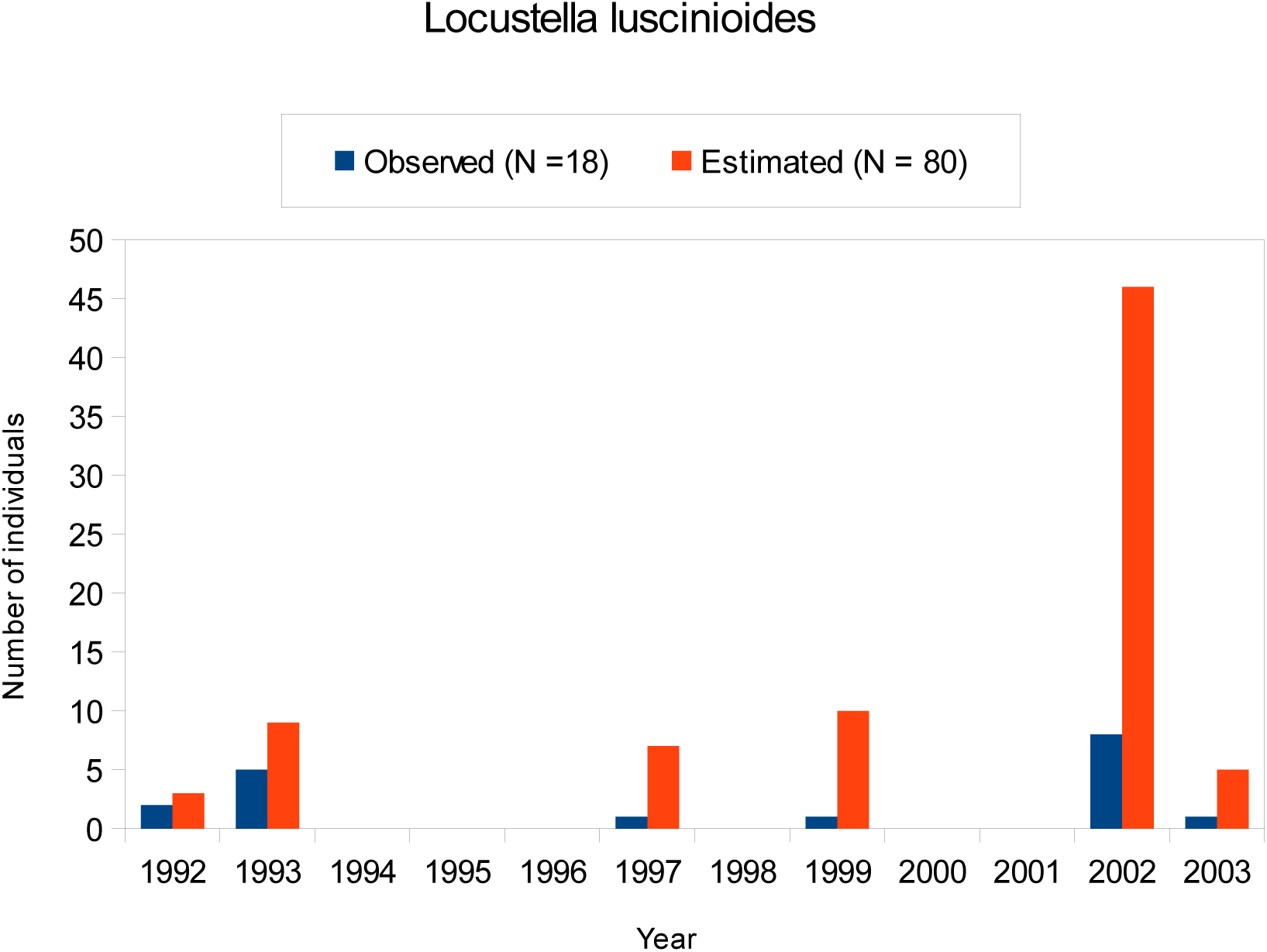
d) Inter-annual phenology of LL in “El Hondo Natural Park”. It shows data from the original sample and the new estimated sample. Original sample (N = 18) and estimated sample(N = 80) matched annually (r = 0.899, t = 6.502, P < 0.001).

### Inter-annual phenology of transaharian migrants in SE Iberia

The original sample did not meet normality annually (Shapiro-Wilk test; LL = 0.669, P = 0.0005; ASCH = 0.815, P = 0.014). The annual sampling effort decreased significantly from 1992 to 2003 (r = - 0.686, t = −2.980, P = 0.014) and was more related to the LL (r = 0.400, P = 0.157) than ASCH (r = 0.299, P = 0.344) abundances. The estimated number of individuals per year was similar (t = 0.092, P = 0.928), andt the sample’s dispersal was similar (LL: CV = 1.944; ASCH: CV = 1.433; Chi-squared: 0.773, P = 0.773) (Figure 2 c, d). The annual estimated frequency distribution was unimodal for LL (Mode 1 = 2001.78) and ASCH (Mode 1 = 1998.96). There was a slight not significant trend towards frequencies’ increases of LL with year (r = 0.09, t = 0.283, P = 0.779) and decreases for ASCH (r = - 0.163, t = −0.523, P = 0.6121). The distribution peak for LL was delayed 3 years in comparison to the ASCH (Median LL= 2002, Median ASCH = 1999; Wilcoxon Test = 4257, P < 0.0001) (Figure 2).

### Climate, phenology and morphological traits

The GLM models selected that the LL intra-annual phenology of Savi’s warblers exhibited a significant positive correlation with intra-annual temperatures, explaining approximately a half of the phenology (Table 3). No morphological trait could be identified as an explanatory factor for these trends, as no significant correlations with abunadances were detected.

## Discussion

The climate dynamics of the Mediterranean are governed by two weather currents (Xoplaki 2002, Lionello *et al*. 2006): the North Atlantic Oscillation (NAO), which influences the weather in the West, and the El Niño Southern Oscillation (ENSO), which influences the weather at Easter. Central Mediterranean areas are mixed by the African monsoon and Sahel rainfall in a phenomenon called the Scandinavian pattern (SCA) that leads to drier areas in SE Iberia and Mediterranean Levant (Bueh and Nakamura 2007, Wang and Tan 2020). This study shows that the local climatic scenario is different from future scenarios, proceeding at least with a microclimatic window explained by a large climatic variability in Iberia (Barrera-Escoda and Cunillera 2011). This could alter the relationships between climate and phenology of these migrants with the rest of Iberia.

The mismatched phenology of LL in relation to ASCH the study area is comparable with the current delayed autumn phenology in space and time as the autumn progresses due to global warming, so the migration pattern of both species is complex (Miholcsa *et al*. 2009) and is positively associated with mean monthly temperatures as expected in central Europe due to climate change (Miholcsa *et al*. 2009).

This study also reveals that LL’s phenology is not associated with the enlargement of structural body size and body condition as the season and year progresses, so the morphological responses of this species to the phenology are not adaptive (Barret 2017) and the increasing population levels of LL are consistent with the population’s models of equilibrium in the Eurasian Reed warbler *Acrocephalus scirpaceus* (Saetre *et al*. 2017) probably because phylogeography considers Iberian LL with stable isolated populations from the last glaciation (Neto *et al*. 2012) and immigration and emigration processes are recent in time, and each time lesser.

Recent evidence suggests that climatic responses to morphology in Boreal birds are maladaptive because implicates shortening of body size polewards (Bosco *et al*. 2022) and that long-term changes in flight-related morphology may affect migration across Europe also in a maladaptive form (e.g. shortening of wing-shape instead of increasing in a long-distance migratory bird in Iberia, Remacha *et al*. 2020).

The positive association of intraphenology of LL with involves different feeding and foraging behaviors and ecomorphological constraints wich affect both warblers. The longer tarsus and feet of LL that indicate a walking motion able to forage in drier and less flooded soils than ASCH (Leisler 1975, Tellería *et al*. 2013 and references therein) which means that LL feeds over more predictable and higher thermo-dependent prey much more abundant in heterogeneous habitats (i.e.halophytic scrub reed-interspersed) and the stay sites could be shorter in space and longer on time than ASCH (e.g. Xeresa marshlands (Valencia: 39°00’N 00°13’W) in a 90 km northern direction which keeps a huge breeding population of LL and Albufera of Valencia (Valencia, 39°17’N 00°20’N) to 153 km in a northeast direction which maintains a median breeding population and with similar habitat features (Aguilella and Álvaro 2006). Since LL didn’t breed in the study period, competition for nesting sites could not account as a limiting factor (Zimin *et al*. 2023) but feeding availability could be dependent on different foraging habits among both warblers during migration. Following Neto et al (2010) feeding behavior of LL is typical of “*foraging-time limited*” migrants because they are limited by food availability due that fuel deposition rate in relation to lean body mass is lower in adults compared with juveniles which may be associated with high intraspecific competition. Since in this study most sample was composed by juveniles, the intraspecific competition should be very weak so the food sources would be very abundant and more free disposable and they regularly would reach the daily metabolisable energy intake and the fuel deposition rate should be slow and low (see mean fat scores of LL, table 1) so at this SE Iberian site they may not to adopt the earlier strategy, limited by food availability (Lindström 2003) but rather the would behave as “*metabolically-limited migrants*” (Lindström 1991).

In ASCH the interannual phenology decreases not significantly with increases of rainfall so is indicative of a declining population on time that is in agreement with the declining populations of transharian migrants in breeding areas (Saino *et al*. 2011, Moller *et al*. 2008). In ASCH body mass decreases in autumn in central Europe (Kovácks *et al*. 2012) and increases from first to last study years whereas in study area anything significant response is found with morphological traits, which suggest that morphological variation processes in this species are not adaptive in SE Iberia. In France, Péron et al (2007) find a positive relationship among stop-over duration and body mass gain in ASCH associated with increases of autumn temperatures.

Multiple factors, such as weather conditions at stopover sites, can affect the phenology of different transaharian migrants (Haest *et al*. 2018, 2020). In ASCH, survival and weather in the Sahel may determine breeding conditions (Peach *et al*. 1991, Tøttrup *et al*. 2012) similarly to other passerines (Boano *et al*. 2004) in which body mass regulation in those areas depends for example on habitat selection (Vafidis *et al*. 2014). The autumn migration of ASCH is negatively related to spring temperatures in Mediterranean areas (Péron *et al*. 2007), and this is consistent with the exponential decrease in annual phenological abundances relative to rainfall for this species in this study. The lack of more significant relationships could be due to the lack of enough data to reach statistical significance.

## Conclusions

Both long-distance migrants are able to adjust their phenology at stopover sites depending on various factors (Gordo and Sanz 2008, Tøttrup *et al*. 2008, Halkka *et al*. 2011, Remisiewicz and Underhill 2022) and depending on a complexity of short-long scale climatic variations (MacMynowski and Root 2007) that may affect the lack of specific-species food (Nartshuk 1996) due to disequilibrium vegetation dynamics and a time shift (e.g. advance) in biomass production enhanced by global warming (Svenning and Sandel 2013; Péron *et al*. 2016). This is study is in agreement with the mismatch hypothesiss of declining migants at the breeding areas due to climatic change ( Jones & Cresswell 2010). Conservation issues of both Afro-tropical migrants need advances in the knowledge to develop and implement effective conservation actions combined with species monitoring and joint adaptive management across the flyway (Vickery *et al*. 2023).

